# Switching Metazoan Fatty Acid Synthase Between Reducing and Non-reducing Elongation Mode via Programming of the Ketoreductase Domain

**DOI:** 10.1101/2025.11.14.688466

**Authors:** Damian L. Ludig, André Herber, Martin Grininger

## Abstract

Polyketides constitute a large class of natural products with important biological activities and applications such as antibiotics, antitumor agents, pesticides, and pigments. Their biosynthesis is catalyzed by polyketide synthases (PKSs) which are multi-domain enzymes evolutionarily related to fatty acid synthases (FASs). Despite their close homology in structure and the chemistry they perform, FASs and PKSs differ fundamentally in their catalytic programming: FASs run fully reducing elongation reactions to yield saturated fatty acids, while iterative PKSs execute reductions just in selected cycles, generating complex oxidized compounds. In this study, we aimed at engineering the metazoan FAS in its KR domain to switch from fully reducing to a non-reducing mode during chain elongation. Guided by recent insights into KR programming, we incorporated a helix into metazoan FAS, which is found in KRs from iterative PKSs and type II FASs with chain length programming. These FAS variants initially catalyze complete fatty acid cycles but lose the ability of β-keto reduction in later elongation rounds, producing intermediates that spontaneously cyclize to pyrone products. Finally, our study provides valuable insight into the mechanism of KR catalysis identifying another amino acid next to the active tyrosine which is capable for intermediate protonation.

## Introduction

Metazoan fatty acid synthase (mFAS), and iterative polyketide synthases (PKSs) are related enzymes that share a conserved multidomain organization and catalytic logic (**Figure 1a** and **b**).^[1]^ Two recent findings have greatly contributed to understanding their evolutionary relationships: First, the discovery of the animal FAS-like polyketide synthases (AFPKs) bridged the phylogenetic gap between iterative PKSs and mFAS, linking secondary-metabolite biosynthesis with primary lipid metabolism.^[2,3]^ Second, iterative type I PKSs were discovered to be widespread not just in fungi, but also in bacteria, linking iterative PKSs and bacterial modular PKSs through a shared evolutionary ancestry.^[4]^ Thus, programming in iterative synthesis is not an isolated fungal innovation, but a deeply conserved catalytic strategy that provides insight into how complex polyketide structures have evolved.^[5]^

**Figure 1.**
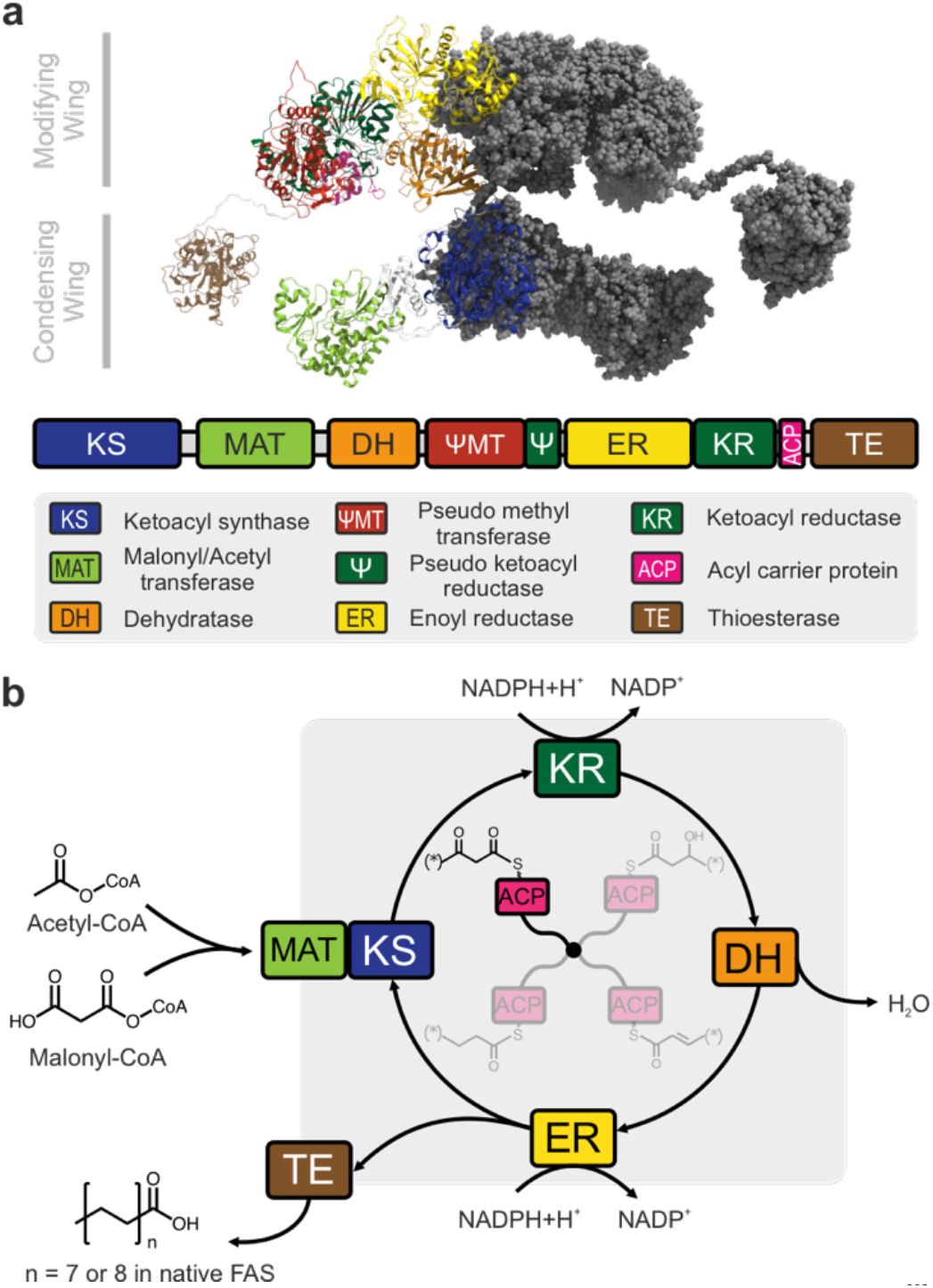
Fatty acid synthase and biosynthesis. (a) Depiction of the dimeric structure of mFAS.^*[6]*^ The domains of one protomer are shown and color-coded as given in the legend. Iterative PKSs share the overall architecture but modulate the modifying domain ensemble. (b) The fatty acid cycle performed by mFAS. During the whole synthesis, intermediates are either bound to a catalytic domain or the acyl carrier protein (ACP) which shuttles substrates and intermediates. The substrates acetyl- and malonyl-CoA are loaded via the MAT. Then, KS forms a C-C bond in a Claisen condensation reaction, resulting in an intermediate which is elongated by two carbon atoms and bears a keto group at the β-position. This β-position can subsequently be processed by three subsequent reactions: i) reduction to the alcohol by the ketoreductase (KR), ii) elimination of the hydroxy group by the dehydratase (DH) and iii) reduction to a saturated chain by the enoylreductase (ER). The resulting saturated acyl chain acts as substrate for the next elongation cycle. Acyl chain of chains length C16 or C18, received after seven or eight cycles respectively, are hydrolyzed from the ACP by the TE and released as free fatty acids.

PKSs are fascinating enzymes due to their ability to catalyze some of the most complex biosynthetic pathways in nature.^[7]^ They are responsible for the biosynthesis of the large and diverse class of polyketides – secondary metabolites that function as antibiotics, antifungals, immunosuppressants, and other bioactive molecules. Beyond a fundamental biochemical interest in this type of proteins, research on PKSs has a large practical perspective, as their engineering offers to expand our therapeutic arsenal. Among PKSs, the modular type proteins are in focus of engineering approaches, because of their apparent programmability. In modular PKSs, each extension unit in the growing polyketide chain is processed by a dedicated module that is used only once during biosynthesis. These modules are arranged in sequence and perform an assembly-line–like synthesis where the order of modules defines the structure of the final product. In contrast, iterative PKSs reuse a single module for multiple rounds of chain extension, making them far less susceptible to engineering. However, the strength in iterative PKSs lies in providing an enzyme-economical yet chemically highly sophisticated biosynthesis. Understanding how these systems control the sequence and outcome of catalytic events, remains one of the central challenges to their rational engineering.^[8]^

FASs have served as tractable models for engineering lipid biosynthesis.^[9–11]^ Their catalytic working mode is strictly uniform to yield saturated products. Considering the decision-making logic of the evolutionarily iterative PKSs, we asked whether the fatty acid biosynthetic regime could be relaxed and mFASs reprogrammed to perform context-dependent catalytic cycles reminiscent of polyketide assembly. Imposing this type of reaction programming onto mFAS would be highly rewarding from an engineering perspective, as mFAS turned out to be a fast^[12]^ and engineerable protein.^[13,14]^

Here, we specifically focus on the ketoreductase (KR) domain, which emerges as a particularly compelling target for programmed iterative synthesis. The KR is a key domain in the programming of the AFPK from *E. chlorotica* (EcPKS1)^[2]^, the tenellin hrPKS/NRPS (TenS)^[15]^ or the desmethylbassianin hrPKS/NRPS (DmbS) (**Figure *2***).^[16]^ The products of EcPKS1, TenS and DmbS feature reduced regions at the ω-end of the chain, while retaining unreduced keto-group(s) in the β (and δ) positions. This observation suggests the presence of a regulatory switch that turns off ketoreduction after initial reductive elongations. As a result, the products appear as arising from the sequential action of a fully reducing FAS-like synthesis and non-reducing PKS-like synthesis, which can also be achieved by a non-reducing FAS (nrFAS) lacking the KR, DH and ER domain (**Figure *2***).^[17]^ In the case of TenS and DmbS, this switch seems to be chain length dependent. Yang *et al*. discovered a helix near the catalytic center of the KR that is responsible for the chain length differences in the TenS/DmbS systems and appears to play a major role in programming of the KR domain.^[18]^

**Figure 2.**
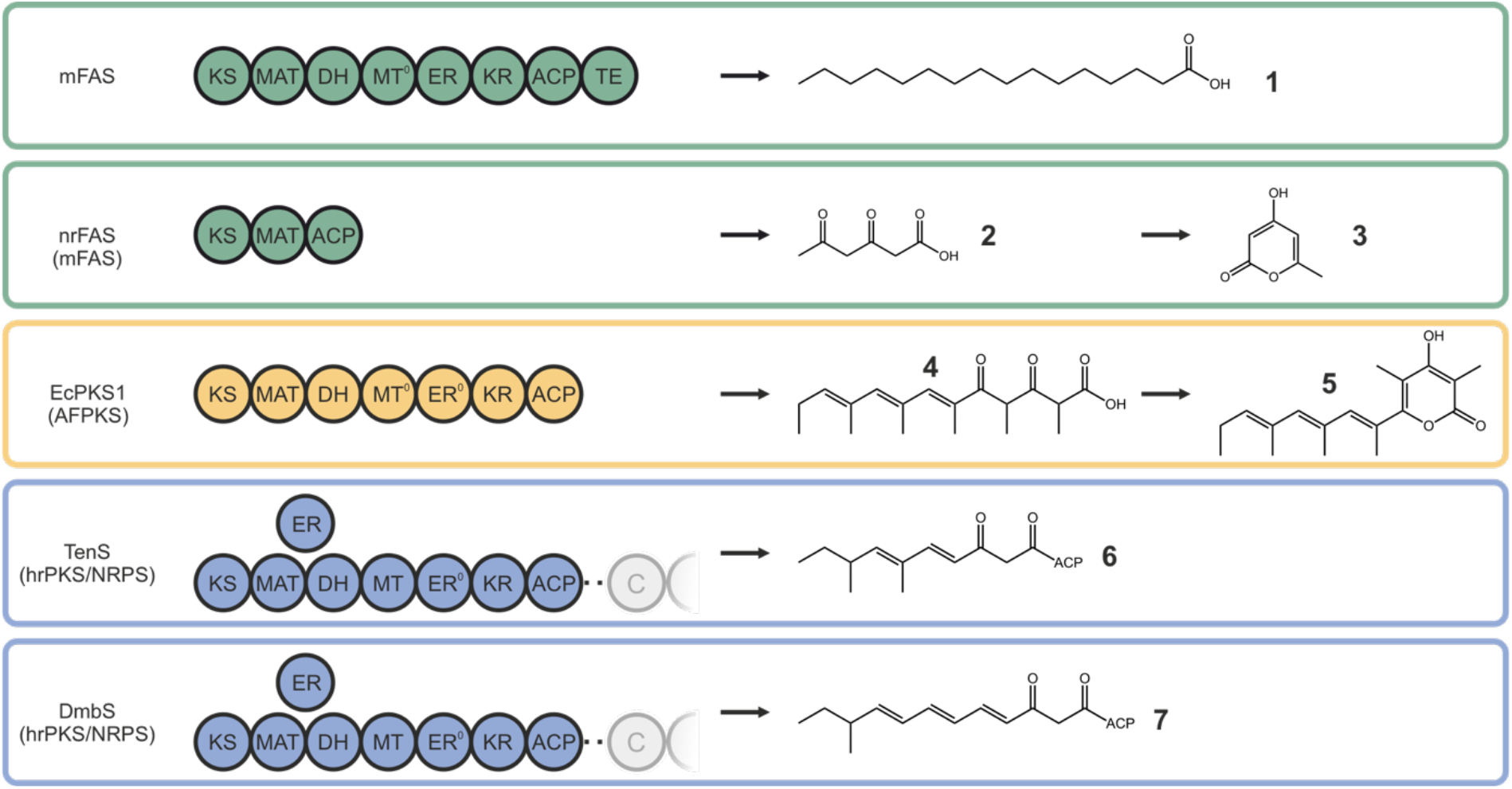
Related FAS/PKS multienzymes and their corresponding products. Domain orders of representatives of mFASs (green), AFPKs (yellow) and hrPKS/NRPS (blue and only PKS part shown). For each protein the corresponding product or ACP-tethered intermediate is shown. mFAS produces palmitic acid (**1**), nrFAS and EcPKS1 produce triketides (**2** and **4**) which form pyrones (**3** and **5**) and TenS and DmbS form ACP-tethered diketides (**6** and **7**).

The pyrone scaffold seen in products **3** and **5** can be found in several natural products with diverse biological activities. For example, 6-alkyl pyrones such as fistupyrone show antifungal activity,^[19]^ while different 2,6-substituted pyrones show antibiotic, antiviral and antiparasitic and anticancer properties.^[20–^

Building on insights from different KR domains, we aimed at reprogramming the KR of mFAS to modulate its reductive behavior. We show that, depending on the specific modification of the KR domain, we can switch between FAS-like elongation and PKS-like elongation at different chain lengths, enabling the production of pyrones with a saturated acyl chain residue. Moreover, this work provides valuable insights into the mechanism of the KR and the roles of the catalytically active amino acid residues.

## Results and Discussion

### Structural Evaluation of KR Programming

To gain insights into the structural determinants of KR programming, we compared the structures of six different KRs (mFAS, EcPKS1, TenS, DmbS and FabG from *H. pylori* and *E. coli*) (**Figure 3** and Figure S1). The structural alignment of the domains showed their similarity in the overall fold and in the active site residues. A major difference was the presence of the substrate binding helix in TenS and DmbS at a position where mFAS contains a flexible loop. Interestingly, FabG features a similar helix to TenS. Specificity data for the *H. pylori* FabG indicate a strong preference for shorter acyl chains,^[23]^ implying a similar way of KR regulation despite its origin from a type II FAS system. In contrast, EcPKS1 lacks the helix structural motif, indicating that the presence of this feature is not the only means of achieving this programming. In our efforts to program mFAS KR, we pursued two different strategies. First, we restricted the space around the catalytic center, next to the active Serine/Tyrosine and the NADPH hydride, by installing sterically demanding amino acids. Second, we replaced the flexible loop with the substrate binding helices from TenS, DmbS, and FabG from *E. coli* and *H. pylori*. (**Figure 3**).

**Figure 3.**
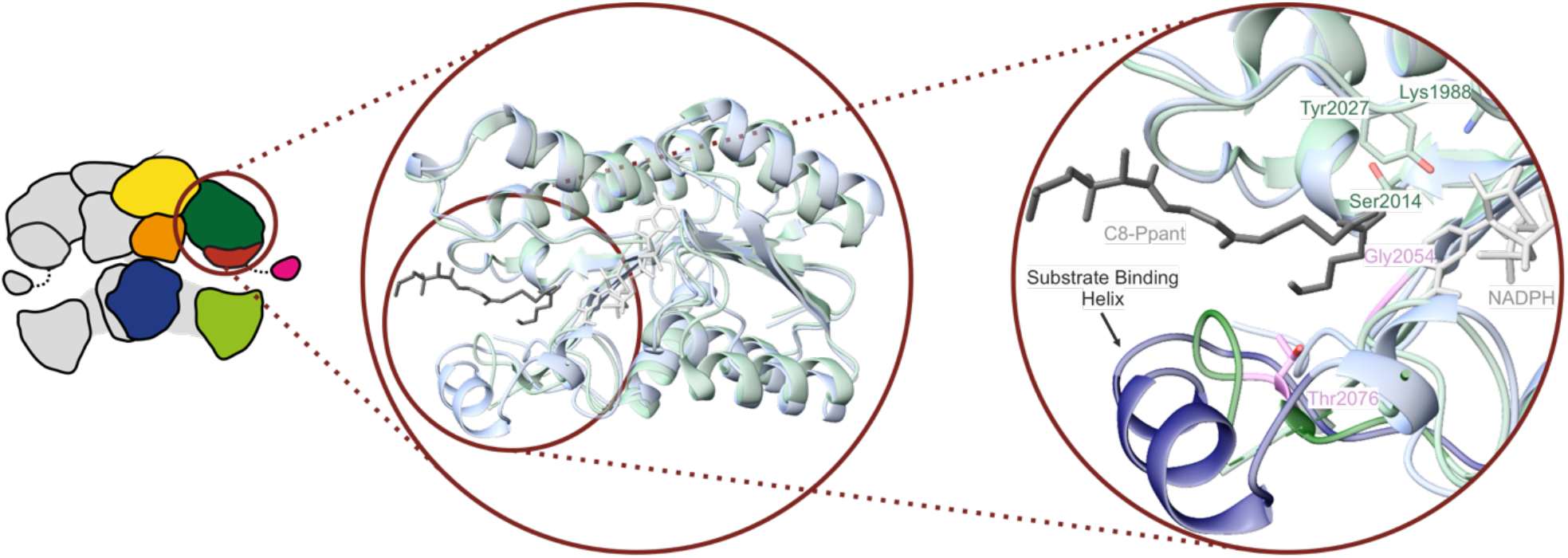
Comparison of mFAS KR and TenS KR. Superposition of the human FAS KR(PDB ID: 8EYI)^*[24]*^ with an Alphafold 3 model of the KR from TenS. Highlighted are the catalytic triad of mFAS KR in green, two positions for point mutations performed in this study in rose, the substrate binding helix of TenS in dark blue and the region which was replaced by the helix in mFAS KR in dark green. The NADPH molecule (light grey) is part of the cryo-EM structure of human FAS KR, the C8-phosphopantetheine (dark grey) is positioned from an overlay with *H. pylori* FabG (PDB ID: 8JFG, Figure S1).^*[23]*^

### Pyrone Formation Activity Assay

mFAS is able to form the pyrone triacetic acid lactone (TAL), when no NADPH is present^[25]^ or as side product in small quantities during standard mFAS synthesis.^[26]^ As observed for EcPKS1^[27]^ or nrFAS,^[17]^ the formation of a pyrone is possible without involving an TE domain, which suggests that the triketide, covalently bound to the ACP, releases spontaneously upon pyrone formation (**Figure 2** and Figure S2).[28]

We harnessed this reaction mode to establish an assay in which an mFAS lacking the TE domain, termed mFASΔTE, is modified in the KR domain. With a KR which has active chain length control, mFASΔTE would first enable reductive elongations to receive fully saturated acyl chain bound to the ACP. At a certain chain length, the KR would then halt reductions and induce non-reductive elongations to the triketide that, once formed, releases spontaneously as pyrone.^[29]^ The pyrone formation can be monitored photometrically, as the absorption at 285 nm is directly proportional to the amount of pyrones formed. We used mFASΔTE as negative control, since it should not produce pyrones in significant amounts under the assay conditions, and nrFAS (KS-MAT-ACP), producing TAL, as positive control.^[17]^ We further constructed two different KR knockouts following the design for 6-deoxyerythronolide B synthase KR6 reported earlier (S2014A and Y2027F, mouse FAS numbering). ^[30]^ These residues stabilize the acyl-substrate and mediate the proton transfer after the nucleophilic attack of the hydride (Figure S3).^[31–33]^

The pyrone formation assays revealed large differences in the effectiveness of our approaches (**Figure 4**). Variants of mFASΔTE, harboring knockout mutations S2014A, Y2027F and S2014A/Y2027F, produced pyrones at different rates. The single point mutations in KR, designed to create steric hindrance, performed differently. While G2054F showed some pyrone production, T2076F apparently failed to disrupt the reductive fatty acid cycle. Finally, most promising results were obtained for the constructs in which the substrate binding helices were inserted. Notably, all four of these constructs were produced in good purity, which is remarkable considering that the inserted substrate binding helices differ between 75% and 94% from the mFAS sequence (Figure S 4-6). All these constructs displayed pyrone production at slightly differing rates of pyrone production. The constructs with helices from FabG exhibited higher pyrone production rates than those with the helices from TenS and DmbS, although their overall activity remained only about half of the activity of nrFAS producing TAL.

**Figure 4.**
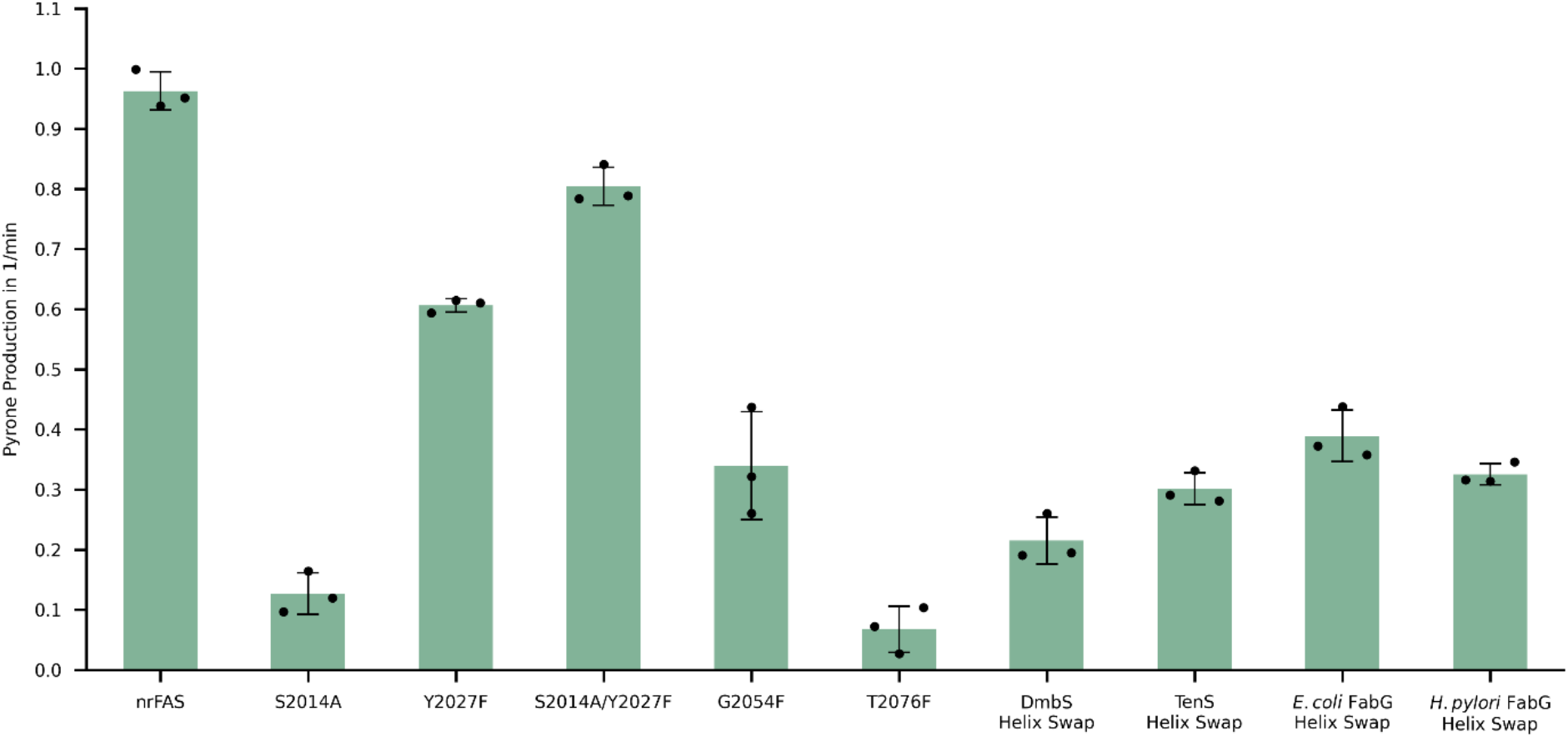
Pyrone formation activity of the KR mutant variants. Pyrone production rate of mFAS mutants monitored by the absorption at 285 nm. Data were collected in biological triplicates, each with three technical replicates. Individual biological replicates are shown as datapoints, the bars represent the means and error bars refer to the standard deviation.

### Product Assay to Determine Chain Length Profile

Building on the outcomes from the pyrone formation activity assay, we next analyzed the constructs regarding the specific pyrones generated. Depending on the type of KR modification, we expected different outcomes: i) KR knockout variants of mFASΔTE were expected to completely prevent reduction, ultimately leading to the production of TAL.^[25]^ ii) Binding site mutants in which small amino acids were replaced by bulkier residues were expected to overall hamper KR activity, leading to an overall broad distribution of pyrones at low concentrations. iii) Helix-insertion variants were expected to most effectively realize the chain length dependent switch, enabling the production of pyrones with an acyl chain.

The binding site mutants showed overall poor switching properties, producing only small amounts of pyrones with up to two reductive elongations (**Figure 5a**). This indicates that they hamper the KR activity, but neither at a significant level nor chain length dependent. In contrast, for the substrate helix constructs, we detected pyrones resulting from up to four reducing elongations before formation of the triketide, although all have propionyl-pyrone as main product. Thus, these variants usually perform one reductive cycle before switched to unreduced elongation. The knockout mutants produced unexpected results, as both the S2014A and Y2027F mutants retained reductive elongation activity. Only when both active residues were mutated simultaneously, the KR was inactive and mFASΔTE exclusively produced TAL. Our data show that the two residues do not contribute to catalysis equally, as mutation Y2027F led to fewer reductive elongations than S2014A. This indicates that Y2027 plays the more critical catalytic role, consistent with its established function as the residue responsible for the protonation of the reaction intermediate (FigS3).^[32–34]^ However, in absence of Y2027, S2014 seems capable of partial compensation, likely by mediating proton transfer through a suitable positioning in the active site (**Figure 5b**). The main role of S2014 appears to lie in aligning the substrate within the active site, a function that Y2027 still can partially maintain in the S2014A mutant.

**Figure 5.**
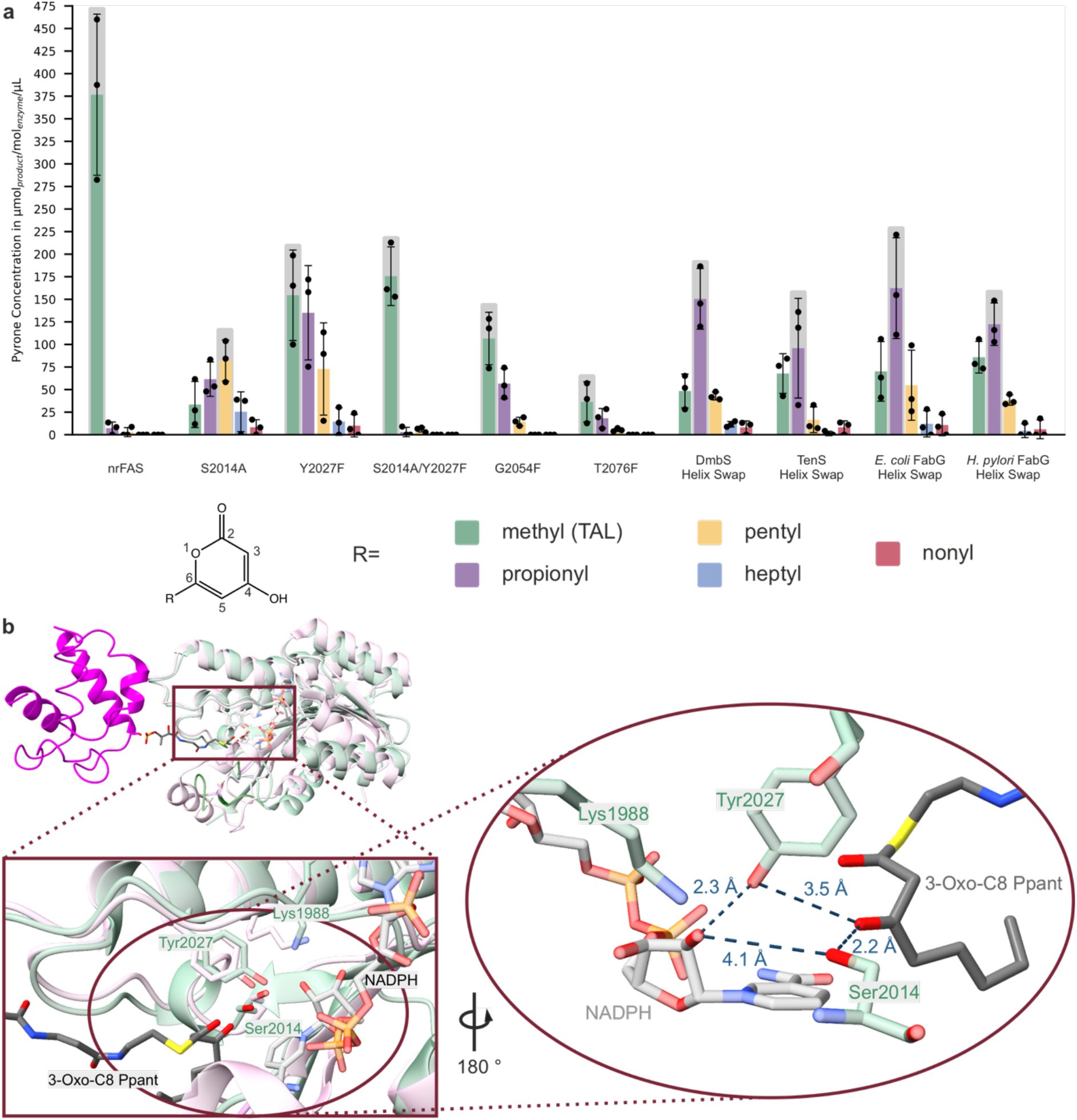
Production of pyrones with different acyl chains on position 6. (a) Product profiles of mFAS mutants analyzed by LC-MS measurements. The major product is highlighted by a grey box. Data were acquired from three independent biological replicates, each measured as three technical replicates. Dots represent individual biological replicates, the bars represent the means and error bars indicate the standard deviation. (b) Overlay of the active sites and position of the ACP-bound 3-Oxo substrate in the active site. A further zoom shows the distances of the catalytic serine and tyrosine to the keto-oxygen and the ribose hydroxygroup, which are the two direct partners in the proton transfer during the reduction. mFAS and NADPH (PDB ID: 8EYI)^*[24]*^, *E. coli* FabG and ACP-bound substrate (PDB ID: 1Q7B)^*[31]*^

Overall, the substrate helix swap constructs displayed a narrower substrate distribution of pyrones compared to knockout and binding site mutants. This is reflected by higher relative abundances of the predominant pyrone chain length over all detected pyrones: 0.39 and 0.40 for S2014G and Y2027F, respectively and 0.58 (DmbS), 0.51 (TenS), 0.52 (*E. coli* FabG) and 0.48 (*H. pylori* FabG). These results demonstrate that the substrate helix adds a layer of substrate specificity onto the mFAS KR, enabling the switch of mFAS from reducing to non-reducing synthesis at defined chain acyl length. We propose that, at a certain chain length, the β-ketoacyl intermediate tethered to the ACP becomes sterically hindered in its interaction with the KR due to the presence of the substrate-binding helix and is then steered towards KS-mediated elongation of the unreduced intermediate. Thus, the relative catalytic efficiencies of the KR and KS for β-ketoacyl chains at a given chain length determine product output.

Likewise, the kinetic interplay of the KS and the TE domains has recently enabled fatty acid chain length control in mFAS.^[13,35]^

## Conclusion

In this work, we investigated the possibilities of switching mFAS between reducing fatty acid and non-reducing polyketide synthesis through the engineering of the KR domain. Via introduction of a substrate binding helix found in some PKS and type II FAS KRs, we achieved the programming of the mFAS KR. In the engineered mFAS, the chain length dependent catalytic efficiencies of the modified KR and KS governed the transition from reductive to non-reductive processing. Specifically, induced by the substrate helix, the KR became kinetically outcompeted by the KS, such that after initial reducing elongation(s), elongation proceeded without reduction, ultimately yielding the triketide that is released as a pyrone. This engineering led, to the best of our knowledge, to the first programmed mFAS KR. The resulting pyrones can serve as direct precursors for the synthesis of more elaborate 3,6 substituted pyrones, an interesting class of natural products.^[36]^

In this work, we demonstrate that chain length programming exhibited by the KR domain in certain iterative PKSs is a transferable feature that can be imposed onto the mFAS framework. This adds an additional layer to the toolbox of mFAS programming and highlights mFAS as a highly engineerable scaffold for programmed biosynthesis.^[12,13]^ From a broader perspective, our results provide evidence that the catalytic decision-making in iterative polyketide synthesis, also referred to as cryptic programming, is modular and transferable, rather than emerging as a complex system-level property. Our findings, therefore, establish a conceptual foundation for the bottom-up programming of biosynthetic pathways within iterative PKSs or in FASs.

## Experimental Section

### Plasmid Transformation

1 μL of the chosen plasmid was added to 50 μL of electrocompetent BAP 1 cell suspension. This mixture was chilled on ice before its transfer to an equally cooled 0.2 cm electroporation cuvette and an electric shock was applied (25 μF, 200 Ω, 2.5 kV). The suspension was immediately removed with the aid of 300 μL prewarmed SOC medium (37 °C) and incubated at 37 °C for 1 h. The cell suspension was centrifuged at 600 rcf for 1.5 min before removal of 300 μL of the supernatant. The cell pellet was resuspended in the remaining supernatant and spread on LB-Agar plates containing 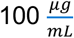Amp before incubation at 37 °C overnight.

### Gene Expression and Protein Purification

20 mL of LB medium containing 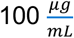Amp per 1 L of main culture were inoculated with five colonies and incubated overnight at 37 °C and 140 rpm. The chosen volume of TB medium containing 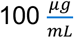 Amp was then inoculated with the pre-culture and incubated at 37 °C and 140 rpm until an optical density (OD) of 0.6–0.8 was reached. The flasks were cooled at 4 °C for 30 min before protein expression was induced by adding 250 μL of 1 M isopropyl β-d-1thiogalactopyranoside (IPTG) per liter main culture. The expression was carried out at 20 °C and 140 rpm overnight. The cells were harvested by centrifugation using an Avanti J20-XP centrifuge with a JLA 8.100 rotor (Beckman Coulter) at 5,000 rcf and 4 °C for 20 min. The supernatant was discarded, and the cell pellet was resuspended in 25 mL Ni-Wash buffer containing a catalytic amount of DNase I and 70 μL of 0.5 M EDTA.

Disruption of the cells was performed at 800 bars using an Amico French Pressure Cell Press (SLM Aminco). The resulting suspension was centrifuged at 50,000 rcf and 4 °C for 1 h using an Avanti J20-XP centrifuge with a JA 25.500 rotor (Beckman Coulter). The supernatant was transferred into a fresh 50 mL falcon tube containing 70 μL of 1 M MgCl2 solution. The crude extract was further purified by affinity chromatography.

The protein of interest was purified from the crude extract by nickel affinity chromatography (3 mL column volume, CV) and following Strep-Tactin ^®^XT chromatography (2 mL CV). Both affinity chromatographic methods include wash and elution phases, each performed with 5 CV of the appropriate buffer (Ni-Wash, Ni-Elution, Strep-Wash, or Strep-Elution, as applicable). The final elution was concentrated to a volume of approximately 1 mL using an Amicon Ultra centrifugal filter unit (Merck). The molecular weight cutoff was chosen appropriately to the protein size. The concentrated protein was stored at −70 °C. All affinity chromatography steps were monitored via SDS-PAGE analysis.

Size exclusion chromatography (SEC) was performed at 4 °C with an ÄKTA Explorer (Cytivia) or an ÄKTA Purifier (Cytivia) equipped with a Superdex 200 Increase 10/300 GL column (Cytiva). All samples were filtered using a 0.22 μm Ultrafree^®^ centrifugal filter (Merck) before application. mFAS samples were incubated at 37 °C for 1 h prior to centrifugation to promote dimerization. Fractions were selected based on the absorbance measured at 280 nm. The combined fractions were concentrated as desired using an Amicon Ultra centrifugal filter unit (Merck) with the appropriate properties. Aliquots were stored at −70 °C.

### Absorption-Based Activity Assay

The activity assay was performed in mFAS assay buffer, the DTT and bovine serum albumin (BSA; final concentration of 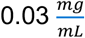) added on the day of the assay. Samples were prepared in a 384-cavity reaction plate (Greiner Bio-One) with a sample volume of 100 μL and measured with a CLARIOstar Plus plate reader (BMG Labtech) at 25 °C. Further settings were set to 240 cycles, 22 flashes per well and cycle, and 15 s cycle time coming to a measurement time of 1 h in which the absorption at 285 nm was monitored. The final concentrations were: 200 µM NADPH, 200 µM Acetyl-CoA, 600 µM Malonyl-CoA and 2 µM enzyme. All enzymes analyzed were purified via SEC. Blank measurements did not include any CoA-esters and were measured using the nrFAS. Samples were prepared in technical triplicates. A calibration curve prepared with TAL was used for all constructs.

### Product Quantification by Liquid Chromatography-Mass Spectrometry

The product assay was performed in mFAS assay buffer, the DTT and bovine serum albumin (BSA; final concentration of 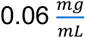) added on the day of the assay. The reaction mixtures with a volume of 150 µL were prepared in 1.5 mL Eppendorf tubes. The final concentrations were: 1400 µM NADPH, 133 µM Acetyl-CoA, 400 µM Malonyl-CoA and 2 µM enzyme. All enzymes analyzed were only purified via affinity chromatographic methods. Samples were kept at room temperature in the dark overnight. Subsequently, 15 μL of 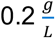 aqueous olivetol was added as the internal standard. 100 μL acetate buffer was added for improved phase separation. The extraction was performed 3 times with 200 μL ethyl acetate (≥ 99.7%, Honeywell). The solvent was removed in a SpeedVac Concentrator SVC-100H (Savant) and substituted with 50 μL methanol (HPLC-grade). After 20 min of centrifugation at 20,000 rcf, 40 μL were submitted in an HPLC-vial for liquid chromatography-mass spectrometry (LC-MS) measurement. Samples were measured on HPLC-ESI-MS using the Ultimate 3000 LC (Dionex) system equipped with an Acquity UPLC BEH C18 (2.1 × 50 mm, particle size 1.7 µm, Waters) and connected to an AmaZonX (Bruker).

Data was analyzed using msconvert^[37]^ and MZmine.^[38]^ The TAL derivatives were confirmed by searching for the theoretical m/z for the [M+H]+-ion. The integrals were further statistically processed using a custom python script using Pandas^[39]^ and Matplotlib.^[40]^

## Conflict of Interest

The authors declare no conflict of interest.

## Acknowledgements

Support for this work was received by intramural funds of the Goethe University Frankfurt. We thank R. Cox for discussion of the project and E. Helfrich and his lab for support in using LC-MS instrumentation.

## Notes

### Competing Interest Statement

The authors have declared no competing interest.

