## Supplementary Information for "Switching Metazoan Fatty Acid Synthase Between Reducing and Non-reducing Elongation Mode via Programming of the Ketoreductase Domain"

**Buffers and Media**

**Table S1:** Buffers used in this work.

| Name | Composition | pH |
|---|---|---|
| mFAS assay buffer | 50 mM KPi, 10% glycerol, 5% PEG 400, 1 mM DTT | 7.0 |
| Ni-Elution | 200 mM KCl, 300 mM imidazole, 50 mM $K_3PO_4$, 10% glycerol | 7.0 |
| Ni-Wash | 200 mM KCl, 30 mM imidazole, 50 mM $K_3PO_4$, 10% glycerol | 7.0 |
| Strep-Elution | 250 mM $K_3PO_4$, 1 mM EDTA, 10% glycerol, 50 mM biotin | 7.0 |
| Strep-Wash | 250 mM $K_3PO_4$, 1 mM EDTA, 10% glycerol | 7.0 |
| acetate buffer | 95.2 mM acetic acid, 4.83 mM sodium acetate | |

**Table S2:** Media used in this work.

| Name | Composition | pH |
|---|---|---|
| LB Agar (Carl Roth) | 1.2% (w/v) bacto agar, 1% (w/v) tryptone, 0.5% (w/v) yeast extract, 0.5% (w/v) NaCl | 7.0 |
| LB medium (Carl Roth) | 1% (w/v) tryptone, 0.5% (w/v) yeast extract, 0.5% (w/v) NaCl | 7.0 |
| SOC medium | 2% tryptone, 0.5% yeast extract, 10 mM NaCl, 2.5 mM KCl, 10 mM MgCl2, 10 mM MgSO4, and 20 mM glucose | - |
| TB medium | 1.2% (w/v) tryptone, 2.4% (w/v) yeast extract, 0.5% (v/v) glycerol, 100 mM phosphate buffer pH 7.5 | 7.5 |

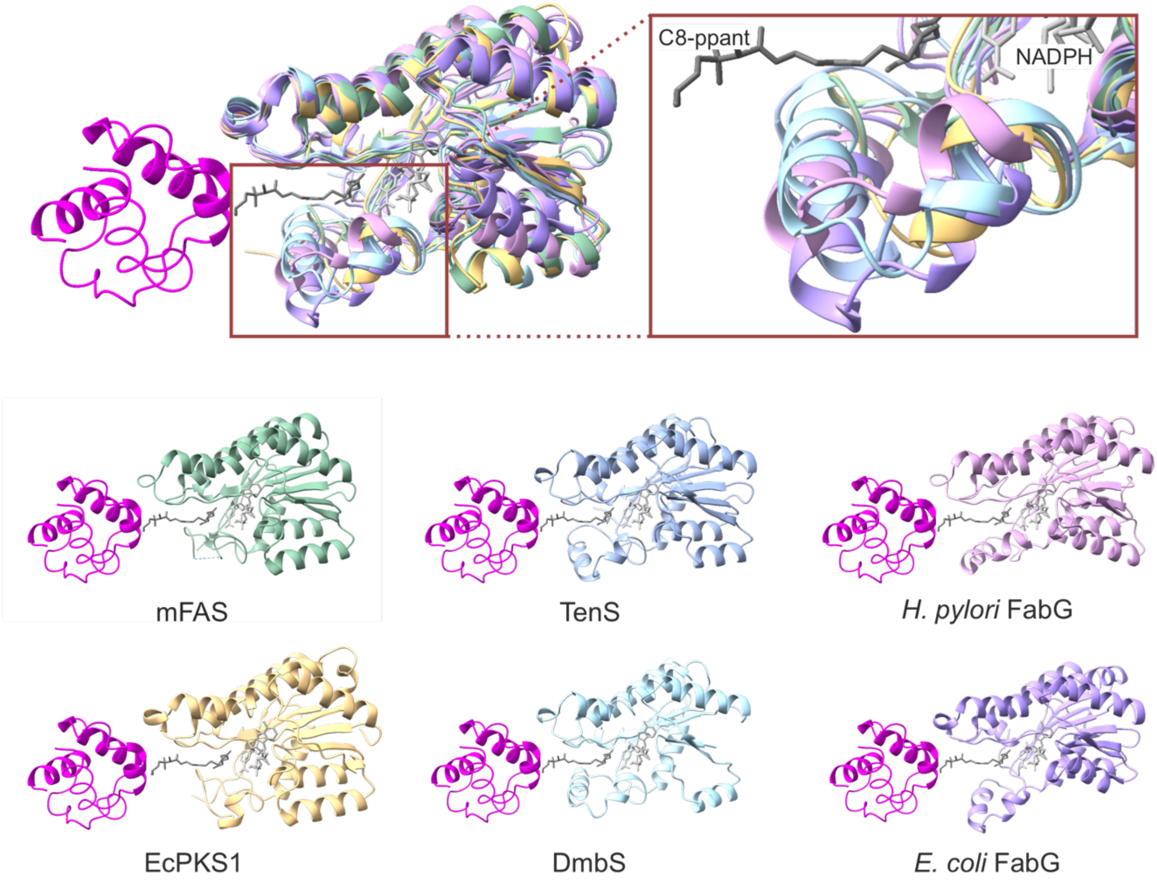


**Figure S1 | Structures of six KRs from different systems and their overlayed binding regions.** mFAS (PDB ID: 8EYI)[14], TenS (Alphafold 3), H. pylori FabG (PDB ID: 8JFG)[13], EcPKS (PDB ID: 9QC9)[27], DmbS (Alphafold 3), E. coli FabG (PDB ID: 1Q7B)[21].

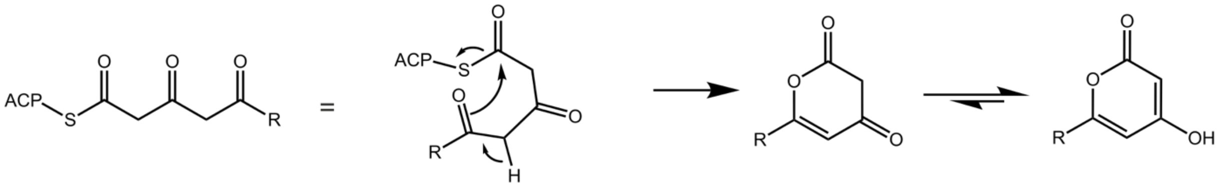


**Figure S2 | Mechanism of the pyrone formation through lactonization of a triketide.** A triketide is released from the ACP by cyclization, followed by a keto-enol tautomerization reaction to leading to a TAL derivative. R represents a variable residue, which, in the case of TAL, is a methyl group.

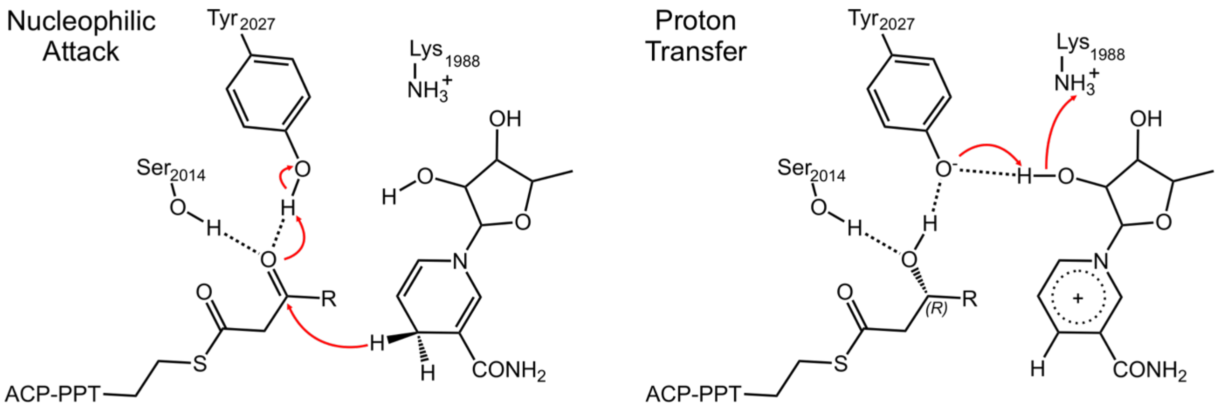


**Figure S3 | Putative mechanism of the KR domain.** Ser2014 and Tyr2027 position the β-keto group for the nucleophilic attack from the hydride of NADPH. When attacked by the hydride, carbonyl oxygen gets protonated by the hydrogen of Tyr2027. Subsequently Tyr2027 gets reprotonated from Lys1988 via proton relay over a hydroxy group of the ribose ring.

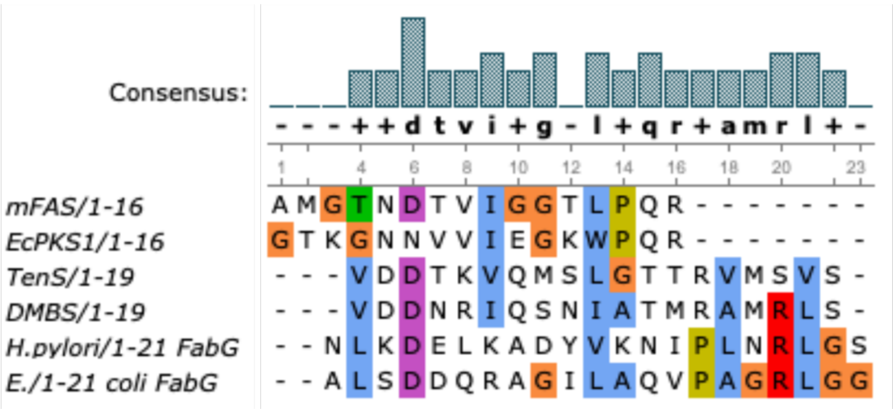


| | mFAS/1-16 | EcPKS1/1-16 | TenS/1-19 | DMBS/1-19 | H.pylori/1-21 FabG | E./1-21 coli FabG |
|---|---|---|---|---|---|---|
| mFAS/1-16 | 100% | 44% | 19% | 13% | 6% | 25% |
| EcPKS1/1-16 | 44% | 100% | 0% | 6% | 0% | 13% |
| TenS/1-19 | 16% | 0% | 100% | 42% | 5% | 11% |
| DMBS/1-19 | 11% | 5% | 42% | 100% | 16% | 26% |
| H.pylori/1-21 FabG | 5% | 0% | 5% | 14% | 100% | 33% |
| E./1-21 coli FabG | 19% | 10% | 10% | 24% | 33% | 100% |

**Legend:** 10% 25% 50% 70% 90%

**Figure S4 | Sequence alignment and similarity matrix of the substrate binding region.** Aligned is the region which builds the substrate binding helices in TenS, DmbS and FabG as well as their counterparts in EcPKS and mFAS.The Alignment was created with Unipro UGENE using the Clustl Algorithm.

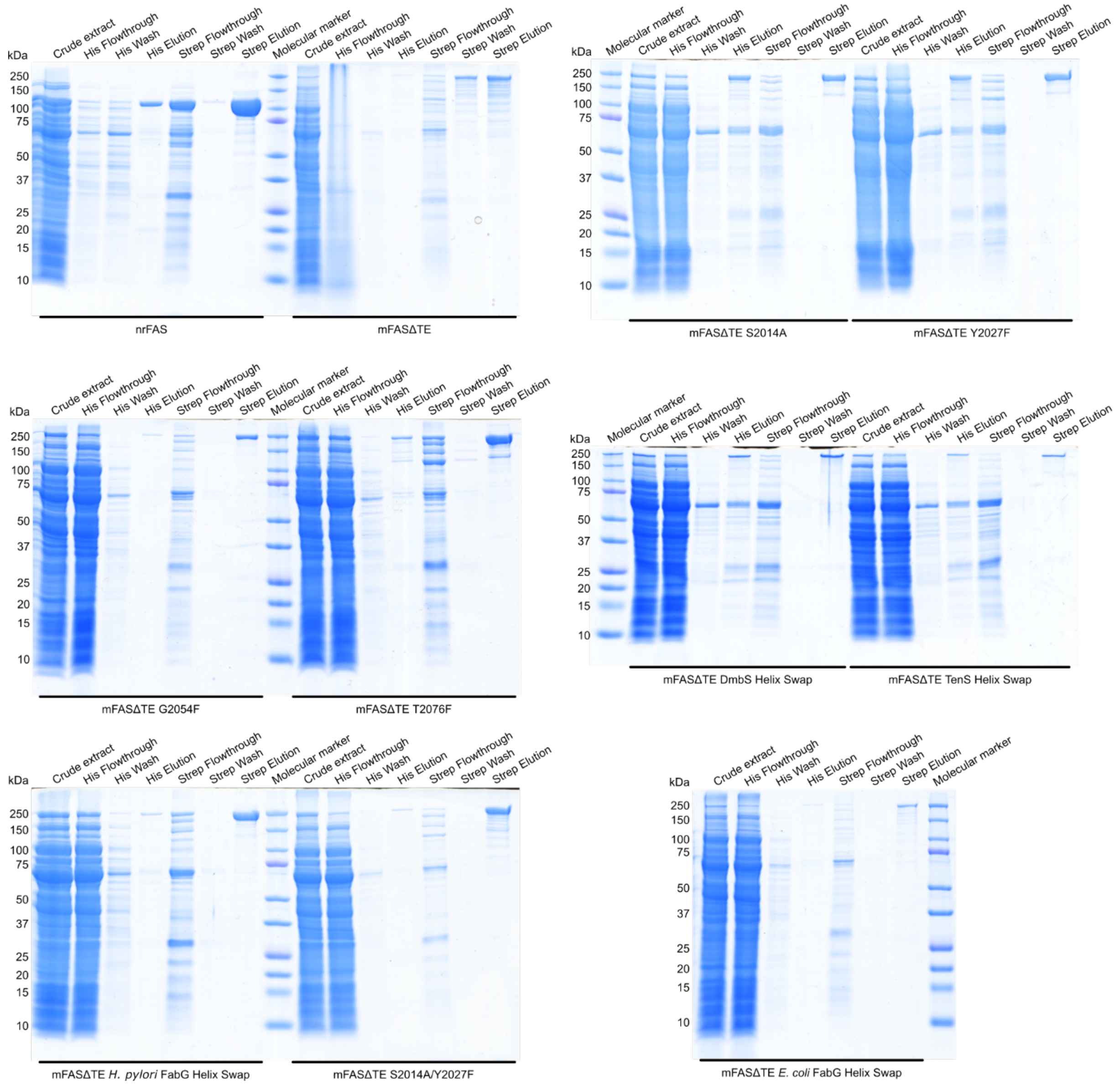


**Figure S5 | Representative SDS-PAGE analysis of the protein purification procedure.** The evident molecular weights of mFAS mutants recombinantly produced in *E. coli* align with the calculated molecular weights of 243–244 kDa and 108 kDa of the mFASΔTE and the nrFAS respectively.

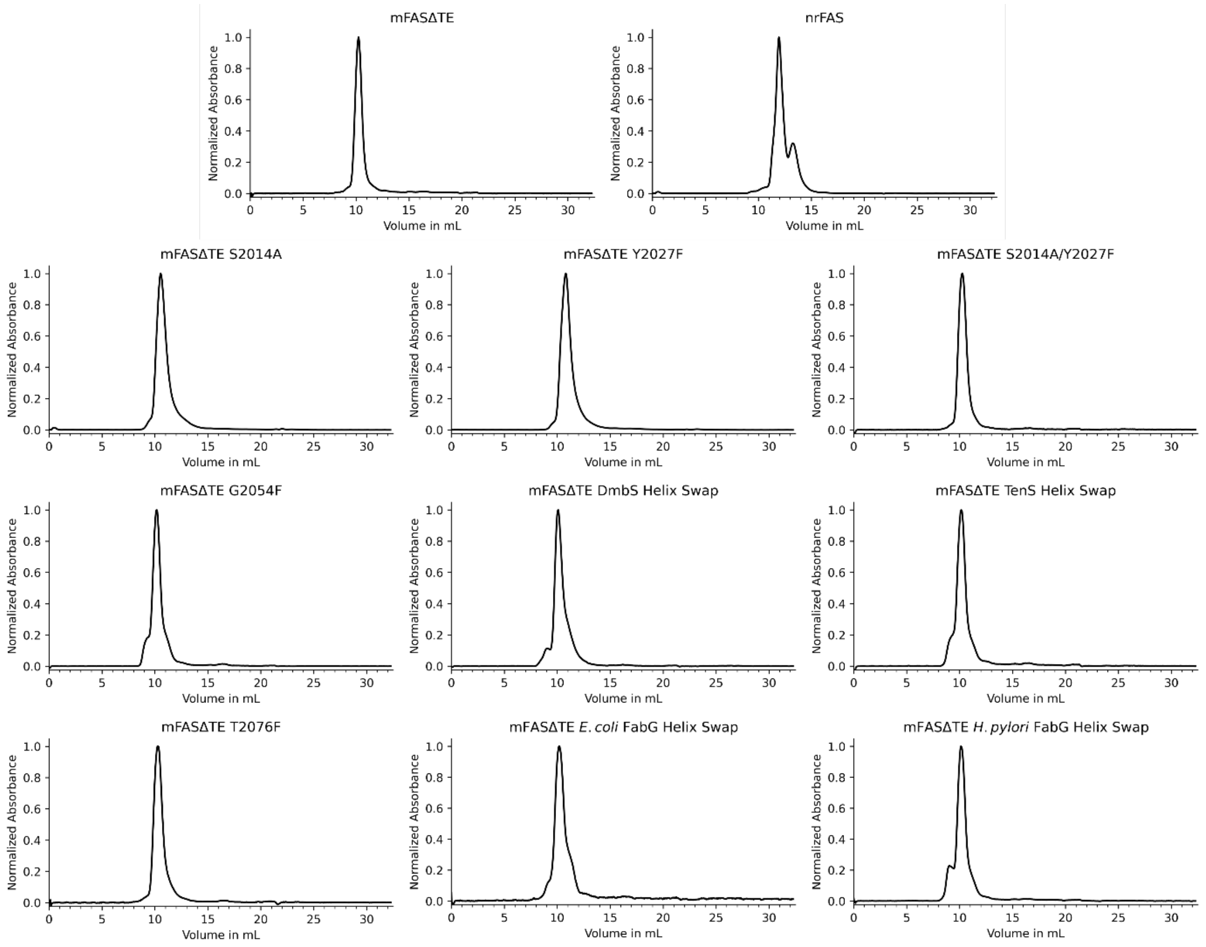


**Figure S6 | Size exclusion chromatogram of all constructs.** The shown normalized chromatograms are representative of all mFAS mutants produced. All mFASΔTE variants show a pronounced peak at 10 to 11 mL elution volume corresponding to the dimeric state of the construct. A following shoulder indicates remaining monomeric protein. A preceding peak/shoulder at 9 mL elution volume points to aggregate. The chromatogram of the nrFAS exhibits the analogues characteristics with a shift +2 mL elution volume. Fractions of the dimeric state were concentrated and used for further experiments.

### Sequence Information

**Table S3 | Primers used in this work.** The fragments for the In-Fusion Cloning containing mutations in the mFAS KR were created using the following primers. Template for all constructs was pAR88 (mFASΔTE).

| Target Mutation | Primer Name | Primer Sequence 5'-3' |
|---|---|---|
| S2014A | PrDL122 | CTACTTTGTGGCCTTCTCCGCAGTAAGCTGCGGGCGTGGTAATG |
| | PrDL123 | CATTACCACGCCCGCAGCTTACTGCGGAGAAGGCCACAAAGTAG |
| Y2027F | PrDL124 | GTAATGCTGGCCAAACTAACTTCGGCTTCGCCAACTCTACCATG |
| | PrDL125 | CATGGTAGAGTTGGCGAAGCCGAAGTTAGTTTGGCCAGCATTAC |
| G2054F | PrDL126 | CCTTGCCGTGCAGTGGTTTGCCATTGGTGACGTGGGC |
| | PrDL127 | GCCCACGTCACCAATGGCAAACCACTGCACGGCAAGG |
| T2076F | PrDL132 | CAATGACACAGTCATCGGAGGTTTCCTGCCTCAGCGCATCTCCTC |
| | PrDL133 | GAGGAGATGCGCTGAGGCAGGAAACCTCCGATGACTGTGTCATTG |
| DmbS Helix Swap | PrDL120 | GCAACATAGCTACCATGCGAGCTATGAGGCTCTCTATCTCCTCCTGCATGGAG |
| | PrDL121 | TGGTAGCTATGTTGCTCTGTATTCTGTTGTCGTCGACTTCCAGGACAATGCCCAC |
| TenS Helix Swap | PrDL118 | TGAGCCTAGGTACCACGCGAGTCATGAGTGTCTCTATCTCCTCCTGCATGGAG |
| | PrDL119 | TGGTACCTAGGCTCATCTGCACCTTGGTGTCGTCAACTTCCAGGACAATGCCCAC |
| *E. coli* FabG Helix Swap | PrDL182 | GTATCCTGGCGCAGGTTCCTGCGGGTCGCCTCGGCGGCATCTCCTCCTGCATGGAG |
| | PrDL183 | CCTGCGCCAGGATACCCGCACGCTGGTCATCGCTCAGCGCTTCCAGGACAATGCCCAC |
| *H. pylori* FabG Helix Swap | PrDL186 | ATTATGTTAAAAACATTCCTTTAAACAGGCTAGGGTCTATCTCCTCCTGCATGGAG |
| | PrDL187 | TGTTTTTAACATAATCCGCTTTGAGTTCGTCTTTCAAATTTTCCAGGACAATGCCCAC |

**Table S4 | Sequence information of the constructs used in this work.** The amino acid and DNA sequences of the constructs pAR88 (mFASΔTE) and pAR127 (nrFAS), as well as for the swapped helices of DmbS, TenS, the *E. coli* FabG, and the *H. pylori* FabG are listed below. The mutated positions are highlighted.

| Construct / Helix | Sequence from start to stop codon / as swapped |
|---|---|
| pAR88 (mFASΔTE, Strep- and His-tagged)<br>Positions 2014 (green), 2027 (cyan), 2054 (magenta), and 2076 (red), as well as the helix swap sequence (yellow) are highlighted | MSAWSHPQFEKGGGSGGGSGGSAWSHPQFEKGAGSEEVVIAGMSGKLPESENLQEFWANLIGGVDMVTDD<br>DRRWKAGLYGLPKRSGKLKDLSKFDASFFGVHPKQAHTMDPQLRLLLEVSYEAIVDGGINPASLRGTNTGV<br>WVGVSGSEASEALSRDPETLLGYSMVGCQRAMMANRLSFFFDFKGPSIALDTACSSSLLALQNAYQAIRSGEC<br>PAALVGGINLLLKPNTSVQFMKLGMLSPDGTCRSFDDSGSGYCRSEAVVAVLLTKKSLARRVYATILNAGTN<br>TDGSKEQGVTFPSGEVQEQLICSLYQPAGLAPESLEYIEAHGTGTKVGDPQELNGITRSLCAFRQAPLLIGSTK<br>SNMGHPEPASGLAALTKVLLSLEHGVWAPNLHFHNPNPEIPALLDGRLQVVDRPLPVRGGNVGINSFGFGGS<br>NVHVILQPNTRQAPAPTAHAALPHLLHASGRTLEAVQDLLEQGRQHSQDLAFVSMLNDIAATPTAAMPFRG<br>YTVLGVEGRVQEVQQVSTNKRPLWFICSGMGTQWRGMGLSLMRLDSFRESILRSDEAVKPLGVKVSDLLLST<br>DERTFDDIVHAFVSLTAIQIALIDLLTSVGLKPDGIIGHSLGEVACGYADGCLSQREAVLAAYWRGQCIKDAHL<br>PPGSMAAVGLSWEECKQRCPAGVVPACHNSEDTVTISGPQAAVNEFVEQLKQEGVFAKEVRTGGLAFHSYF<br>MEGIAPTLLQALKKVIREPRPRSARWLSTSIPEAQWQSSLARTSSAEYNVNNLVSPVLFQEALWHIPEHAVV<br>LEIAPHALLQAVLKRGVKSSCTIIPLMKRDHKDNLEFFLTNLGKVHLTGINVNPNALFPPVEFPAPRGTPLISP<br>HIKWDHSQTWDVPVAEDFPNGSSSSSATVYSIDASPESPDHYLVDHCIDGRVIFPGTGYLCLVWKTLARSLGL<br>SLEETPVVFENVSFHQATILPKTGTVALEVRLLEASHAFEVSDTGNLIVSGKVYLWEDPNSKLFDHPEVPTPP<br>ESASVSRLTQGEVYKELRLRGYDYGPQFQGICEATLEGEQGKLLWKDNWVTFMDTMLQVSILGSSQQSLQLP<br>TRVTAIYIDPATHRQKVYRLKEDTQVADVTTSRCLGITVSGGIHISRLQTTATSRRQQEQLVPTLEKFVFTPH<br>MEAECLSESTALQKELQLCKGLARALQTKATQQGLKAAMLGQEDPPQHGLPRLLAAACQLQLNGNLQLELG<br>EALAQERLLLPEDPLISGLLNSQALKACVDTALENLSTLKMKVAEVLAGEGHLYSRIPALLNTQPMLQLEYTA<br>TDRHPQALKDVQTKLQQHDVAQGQWNPSDPAPSSLGALDLLVCNCALATLGDPALALDNMVAALKEGGFL<br>LVHTVLKGHALGETLACLPSEVQPAPSLLSQEEWESLFSRKALHLVGLKRSFYGTALFLCRRAIPQEKPIFLSV<br>EDTSFQWVDSLKSTLATSSSQPVWLTAMDCPTSGVVGLVNCLRKEPGGHRIRCILLSNLSNTSHAPKLDPGS<br>PELQQVLKHDLVMNVYRDGAWGAFRHFQLEQDKPKEQTAHAFVNVLTRGDLASIRWVSSPLKHTQPSSSG<br>AQLCTVYYASLNFRDIMLATGKLSPDAIPGKWASRDCMLGMEFSGRDRCGRRVMGLVPAEGLATSVLLSSDF<br>LWDVPSSWTLEEAASVPVVYTTAYYSLVVRGRIQRGETVLIHSGSGGVGQAAISIALSLGCRVFTTVGSAEKR<br>AYLQARFPQLDDTSFANSRDTSFEQHVLLHTGGKGVDLVLNSLAEEKLQASVRCLAQHGRFLEIGKFDLSNN<br>HPLGMAIFLKNVTFHGILLDALFEEANDSWREVAALLKAGIRDGVVKPLKCTVFPKAQVEDAFRYMAQGKH<br>IGKVLVQVREEEPEAVLPGAQPTLISAISKTFCPAHKSYIITGGLGGFGLELARWLVLRGAQRLVLTSRSGIRTG<br>YQAKHIREWRRQGIQVLVSTSNVSSLEGARALIAEATKLGPVGGVFNLAMVLRDAMLENQTPELFQDVNKP<br>KYNGTLNLDRATREACPELDYFVAFSSVSCGRGNAGQTNYGFANSTMERICEQRRHDGLPGLAVQWGAIGD<br>VGIVLEAMGTNDTVIGGTLPQRISSCMEVLDLFLNQPHAVLSSFVLAEKKAVAHGDGDTQRDLVKAVAHILGI<br>RDLAGINLDSTLADLGLDSLMGVEVRQILEREHDLVLPMREVRQLTLRKLQEMSSKTDSATDTTLEHHHHH<br>HHH |

| Construct / Helix | Sequence from start to stop codon / as swapped |
| --- | --- |
| pAR88 (mFASΔTE, Strep- and His-tagged)<br>Positions 2014 (green), 2027 (cyan), 2054 (magenta), and 2076 (red), as well as the helix swap sequence (yellow) are highlighted | ATGAGCGCTTGGAGCCATCCACAATTTGAGAAGGGTGGAGGTTCTGGCGGTGGATCGGGAGGTTCAGCGTG<br>GAGCCACCCGCAGTTCGAAAAAGGCGCCGGATCCGAGGAGGTGGTGATAGCCGGTATGTCGGGGAAGTTGC<br>CCGAGTCAGAGAACCTACAGGAGTTCTGGGCCAACCTCATTGGTGGTGTGGACATGGTCACAGATGATGAC<br>AGGAGATGGAAGGCTGGGCTCTATGGATTACCCAAGCGGTCTGGAAAGCTGAAGGATCTCTCCAAGTTCGA<br>CGCCTCCTTTTTTGGGGTCCACCCCAAGCAGGCACACACAATGGACCCCCAGCTTCGGCTGCTGTTGGAAGT<br>CAGCTATGAAGCAATTGTGGATGGAGGTATCAACCCAGCCTCACTCCGAGGAACGAACACTGGCGTCTGGG<br>TGGGTGTGAGTGGTTCAGAGGCATCCGAGGCCCTTAGCAGAGATCCCGAGACGCTTCTGGGCTACAGCATG<br>GTGGGCTGCCAGCGTGCAATGATGGCCAACCGGCTCTCTTTCTTCTTCGACTTCAAAGGACCAAGCATTGCC<br>CTGGACACAGCCTGCTCCTCCAGCTTGCTGGCACTACAGAATGCCTACCAGGCCATCCGTAGTGGGGAATGC<br>CCCGCGGCCCTTGTGGGTGGGATCAACCTGCTCCTGAAGCCGAACACCTCTGTGCAGTTCATGAAGCTGGGC<br>ATGCTCAGCCCGGACGGCACCTGCAGATCCTTTGATGATTCAGGGAGTGGATATTGTCGCTCTGAGGCTGT<br>TGTAGCAGTTCTGCTGACTAAGAAGTCCCTGGCTCGGCGGGTCTATGCCACGATTCTGAATGCCGGCACCAA<br>TACAGATGGCAGCAAGGAGCAAGGTGTAACATTCCCCTCTGGAGAAGTCCAAGAACAACTCATCTGCTCTC<br>TGTATCAGCCAGCTGGTCTGGCCCCGGAGTCGCTTGAGTATATTGAAGCCCATGGCACGGGCACCAAGGTG<br>GGTGACCCCCAGGAACTGAATGGCATTACTCGGTCCCTGTGCGCCTTCCGCCAGGCCCCTCTGTTAATTGGC<br>TCCACCAAATCCAACATGGGACACCCTGAGCCTGCCTCTGGGCTTGCAGCCCTGACCAAGGTGCTGTTATCC<br>CTGGAGCATGGGGTCTGGGCCCCTAACCTGCACTTCCACAACCCCAACCCTGAGATCCCAGCACTTCTTGAT<br>GGGCGGCTGCAGGTGGTCGATAGGCCCCTGCCTGTTCGTGGTGGCAACGTGGGCATCAACTCATTTGGCTTC<br>GGAGGCTCCAATGTTCATGTCATCCTCCAGCCCAACACACGGCAGGCCCCTGCGCCCACTGCACACGCTGCC<br>CTTCCCCATTTGCTGCACGCCAGTGGACGCACCTTAGAGGCAGTGCAGGACCTGCTGGAACAGGGCCGCCAG<br>CACAGCCAGGACCTGGCCTTTGTGAGCATGCTCAATGACATTGCGGCAACCCCTACAGCAGCCATGCCCTTC<br>AGGGGTTACACTGTGCTAGGTGTTGAGGGCCGTGTCCAAGAAGTGCAGCAAGTGTCCACCAACAAGCGCCC<br>ACTCTGGTTCATCTGCTCAGGGATGGGCACGCAGTGGCGCGGGATGGGGCTGAGCCTCATGCGCCTGGACA<br>GCTTCCGTGAGTCTATCCTGCGCTCCGATGAGGCTGTGAAGCCGTTGGGAGTGAAAGTGTCAGATCTGCTG<br>TTGAGCACAGATGAGCGCACCTTTGATGACATCGTGCATGCCTTTGTGAGCCTCACTGCCATCCAGATTGCC<br>CTCATCGACCTACTGACTTCTGTGGGACTGAAACCTGACGGCATCATTGGGCACTCCTTGGGAGAGGTTGCC<br>TGTGGCTATGCAGATGGCTGTCTCTCCCAGAGAGAGGCTGTGCTTGCAGCTTACTGGCGAGGCCAGTGCAT<br>CAAAGATGCCCACCTCCCGCCTGGATCCATGGCAGCTGTTGGTTTGTCCTGGGAGGAATGTAAACAGCGCTG<br>CCCCGCTGGCGTGGTGCCTGCCTGCCACAACTCTGAGGACACCGTGACCATCTCTGGACCTCAGGCTGCAGT<br>GAATGAATTTGTGGAGCAGCTAAAGCAAGAAGGTGTGTTTGCCAAGGAGGTACGAACAGGAGGCCTGGCTT<br>TCCACTCCTACTTCATGGAAGGAATTGCCCCCACATTGCTGCAGGCTCTCAAGAAGGTGATCCGGGAACCAC<br>GGCCGCGCTCGGCTCGATGGCTCAGCACCTCTATCCCTGAGGCCCAGTGGCAGAGCAGCCTGGCCCGCACAT<br>CTTCTGCCGAGTACAATGTCAACAACCTGGTGAGCCCTGTGCTCTTCCAGGAAGCACTGTGGCACATCCCTG<br>AGCATGCCGTGGTGCTGGAGATTGCGCCCCACGCACTGTTGCAGGCTGTCCTGAAGCGAGGCGTGAAGTCC<br>AGCTGCACCATCATTCCCTTGATGAAGAGGGATCATAAAGATAACTTGGAGTTCTTTCTCACCAACCTTGG<br>CAAGGTGCACCTCACAGGCATCAATGTCAACCCTAACGCCTTGTTCCCACCTGTGGAGTTCCCGGCTCCCCG<br>AGGGACTCCTCTCATCTCCCCTCACATCAAGTGGGACCACAGTCAGACTTGGGATGTCCCGGTTGCTGAGGA<br>CTTCCCAAACGGCTCCAGCTCCTCCTCTGCTACAGTCTACAGCATCGACGCCAGTCCTGAGTCGCCCGACCA<br>CTACCTGGTAGACCACTGCATTGACGGCCGGGTCATCTTCCCTGGCACTGGCTACCTGTGCCTGGTGTGGAA<br>GACACTGGCTCGCAGCCTGGGCTTGTCCCTAGAAGAGACCCCTGTGGTATTTGAGAATGTGTCGTTTCATC<br>AGGCCACTATACTACCCAAGACAGGAACCGTGGCGCTGGAGGTGAGGCTGCTAGAGGCCTCCCATGCCTTT<br>GAGGTGTCTGACACTGGCAATCTGATTGTGAGCGGAAAAGTGTACCTGTGGGAAGACCCGAACTCCAAGTT<br>ATTCGACCACCCAGAAGTCCCAACACCCCCTGAGTCTGCATCGGTCTCCCGCCTGACCCAGGGAGAAGTATA<br>CAAGGAGCTGCGGCTGCGTGGCTATGATTATGGCCCTCAGTTCCAGGGCATCTGTGAGGCCACCCTTGAAG<br>GTGAACAAGGCAAGCTGCTCTGGAAAGATAACTGGGTGACCTTCATGGACACAATGCTGCAGGTATCCATT<br>CTGGGTTCTAGCCAGCAGAGTCTACAGCTACCTACCCGTGTGACCGCCATCTATATCGACCCTGCCACCCAC<br>CGTCAGAAGGTGTACAGGCTGAAGGAGGACACTCAAGTGGCTGATGTGACAACGAGCCGCTGTCTGGGCAT<br>AACGGTCTCTGGTGGTATCCACATCTCAAGACTACAGACGACAGCAACCTCACGGCGGCAGCAAGAACAGC<br>TGGTCCCCACCTTGGAAAAGTTCGTTTTCACACCGCACATGGAGGCTGAGTGCCTGTCTGAGAGCACTGCCC<br>TGCAGAAGGAGCTGCAACTGTGCAAGGGTCTGGCACGGGCTCTGCAGACCAAGGCCACCCAGCAAGGGCTG<br>AAGGCGGCAATGCTTGGGCAAGAGGACCCTCCACAGCACGGGCTGCCTCGACTCCTGGCAGCTGCTTGCCAG<br>TTGCAGCTCAACGGGAACCTGCAGCTGGAGCTGGGAGAAGCGCTGGCTCAAGAGAGGCTCCTGCTGCCAGA<br>AGACCCTCTGATCAGTGGCCTCCTCAACTCCCAGGCCCTCAAGGCCTGCGTAGACACAGCCCTGGAGAACTT<br>GTCTACTCTCAAGATGAAGGTGGCAGAGGTGCTGGCTGGAGAAGGCCACTTGTATTCCCGAATCCCGGCAC<br>TGCTCAACACCCAGCCCATGCTACAACTGGAATACACAGCCACCGACCGGCACCCCCAGGCCCTGAAGGATG<br>TTCAGACCAAACTGCAGCAGCATGATGTGGCGCAGGGCCAGTGGAACCCTTCCGACCCTGCGCCCAGCAGCC<br>TGGGTGCCCTTGACCTTCTGGTGTGCAACTGTGCATTAGCCACCCTGGGGGATCCAGCCTTGGCCCTGGACA<br>ACATGGTAGCTGCCCTCAAGGAAGGTGGTTTCCTGCTAGTGCACACAGTGCTCAAAGGACATGCCCTTGGG<br>GAGACCCTGGCCTGCCTACCCTCTGAGGTGCAGCCTGCGCCCAGCCTCCTAAGCCAGGAGGAGTGGGAGAGC<br>CTGTTCTCGAGGAAGGCACTACACCTGGTGGGCCTTAAAAGGTCCTTCTACGGTACTGCGCTGTTCCTGTGC<br>CGGCGAGCCATCCCACAGGAGAAACCTATCTTCCTGTCTGTGGAGGATACCAGCTTCCAGTGGGTGGACTCT<br>CTGAAGAGCACTCTGGCCACGTCCTCCTCCCAGCCTGTGTGGCTAACGGCCATGGACTGCCCCACCTCGGGT<br>GTGGTGGGTTTGGTGAATTGTCTCCGAAAAGAGCCGGGTGGACACCGGATTCGGTGTATCCTGCTGTCCAA<br>CCTCAGCAACACATCTCACGCCCCCAAGTTGGACCCTGGCTCTCCAGAGCTACAGCAGGTGCTAAAGCATGA<br>CCTCGTGATGAACGTGTACCGGGACGGGGCCTGGGGTGCCTTCCGTCACTTCCAGTTAGAGCAGGACAAGCC<br>CAAGGAGCAGACAGCGCATGCCTTTGTAAACGTCCTCACCCGAGGGGACCTCGCCTCCATCCGCTGGGTCTC<br>CTCCCCCCTGAAGCACACGCAGCCCTCGAGCTCAGGAGCACAGCTCTGCACTGTCTACTACGCCTCACTGAA<br>CTTCCGAGACATCATGCTGGCCACGGGCAAGCTGTCCCCTGATGCCATTCCAGGTAAATGGGCCAGCCGAGA<br>CTGCATGCTCGGCATGGAGTTCTCAGGCCGGGATAGGTGTGGCCGGCGTGTGATGGGGCTGGTTCCTGCAG<br>AAGGCCTGGCCACCTCAGTCCTGCTATCATCTGACTTCCTCTGGGATGTACCCTCCAGCTGGACCCTGGAGG<br>AGGCGGCCTCTGTGCCCGTCGTCTATACCACTGCTTACTACTCGTTAGTGGTTCGCGGGCGCATCCAGCGTG<br>GGGAGACCGTGCTCATCCACTCAGGTTCAGGTGGTGTGGGCCAAGCGGCCATTTCCATTGCCCTCAGTCTGG<br>GCTGCCGCGTCTTCACCACTGTGGGCTCTGCAGAGAAGCGAGCATACCTCCAGGCCAGGTTCCCTCAGCTTG<br>ATGACACCAGCTTTGCCAACTCGAGGGACACATCATTTGAGCAGCACGTGTTACTGCACACAGGTGGCAAA |

| Construct / Helix | Sequence from start to stop codon / as swapped |
|---|---|
| | GGGGTCGACCTGGTCCTCAACTCACTGGCAGAAGAGAAGCTGCAGGCCAGTGTGCGGTGCTTGGCTCAGCA<br>TGGTCGCTTCTTAGAGATTGGCAAATTTGATCTTTCTAACAACCACCCTCTGGGCATGGCTATCTTCTTGAA<br>GAACGTCACTTTCCATGGGATCCTGCTGGACGCCCTTTTTGAGGAGGCCAATGACAGCTGGCGGGAGGTGG<br>CGGCACTCCTGAAGGCTGGCATTCGTGATGGAGTCGTGAAGCCCCTCAAGTGCACAGTGTTTCCCAAGGCCC<br>AGGTGGAAGATGCCTTCCGCTACATGGCTCAGGGGAAACACATTGGCAAAGTCCTTGTCCAGGTACGGGAG<br>GAGGAGCCTGAGGCTGTGCTGCCAGGGGCTCAGCCCACCCTGATTTCTGCCATCTCCAAGACCTTCTGCCCA<br>GCCCATAAGAGTTACATCATCACTGGTGGCCTAGGTGGCTTTGGCCTGGAGCTGGCCCGGTGGCTCGTGCTT<br>CGCGGAGCCCAGAGGCTTGTGCTGACTTCCCGATCTGGAATCCGCACCGGCTACCAAGCCAAGCACATTCGG<br>GAGTGGAGACGCCAGGGCATCCAAGTGCTCGTGTCAACAAGCAACGTGAGCTCACTGGAGGGGGCCCGTGC<br>TCTCATCGCCGAAGCCACAAAGCTGGGGCCCGTTGGGGGTGTCTTCAACCTGGCCATGGTTTTGAGGGATGC<br>CATGCTGGAGAACCAGACCCCAGAGCTCTTCCAGGATGTCAACAAGCCCAAATACAATGGCACCCTGAACCT<br>TGACAGGGCAACCCGGGAAGCCTGCCCTGAGCTGGACTACTTTGTGGCCTTCTCCTCTGTAAGCTGCGGGCG<br>TGGTAATGCTGGCCAAACTAACTACGGCTTCGCCAACTCTACCATGGAGCGTATATGTGAACAGCGCAGGC<br>ACGATGGCCTCCCAGGCCTTGCCGTGCAGTGGGGTGCCATTGGTGACGTGGGCATTGTCCTGGAAGCGATG<br>GGCACCAATGACACAGTCATCGGAGGTACGCTGCCTCAGCGCATCTCCTCCTGCATGGAGGTACTGGACCTC<br>TTCCTGAATCAGCCCCACGCAGTCCTGAGCAGCTTTGTGCTGGCAGAGAAGAAAGCTGTGGCCCATGGGGA<br>CGGGGACACCCAGAGGGATCTGGTGAAAGCTGTAGCACACATCCTAGGCATCCGAGACCTCGCAGGTATTA<br>ACCTGGACAGCACGCTGGCAGACCTCGGCCTGGACTCGCTCATGGGTGTGGAAGTTCGTCAGATCCTGGAAC<br>GAGAACACGATCTGGTGCTGCCCATGCGTGAGGTGCGGCAGCTCACGCTGCGGAAACTTCAGGAAATGTCC<br>TCCAAGACTGACTCGGCTACTGACACGACACTCGAGCATCATCACCACCACCACCACCAC |
| pAR127 (nrFAS, Strep- and His-tagged) | MSAWSHPQFEKGGGSGGGSGGSAWSHPQFEKGAGSEEVVIAGMSGKLPESENLQEFWANLIGGVDMVTDD<br>DRRWKAGLYGLPKRSGKLKDLSKFDASFFGVHPKQAHTMDPQLRLLLEVSYEAIVDGGINPASLRGTNTGV<br>WVGVSGSEASEALSRDPETLLGYSMVGCQRAMMANRLSFFFDFKGPSIALDTACSSSLLALQNAYQAIRSGEC<br>PAALVGGINLLLKPNTSVQFMKLGMLSPDGTCRSFDDSGSGYCRSEAVVAVLLTKKSLARRVYATILNAGTN<br>TDGSKEQGVTFPSGEVQEQLICSLYQPAGLAPESLEYIEAHGTGTKVGDPQELNGITRSLCAFRQAPLLIGSTK<br>SNMGHPEPASGLAALTKVLLSLEHGVWAPNLHFHNPNPEIPALLDGRLQVVDRPLPVRGGNVGINSFGFGGS<br>NVHVILQPNTRQAPAPTAHAALPHLLHASGRTLEAVQDLLEQGRQHSQDLAFVSMLNDIAATPTAAMPFRG<br>YTVLGVEGRVQEVQQVSTNKRPLWFICSGMGTQWRGMGLSLMRLDSFRESILRSDEAVKPLGVKVSDLLLST<br>DERTFDDIVHAFVSLTAIQIALIDLLTSVGLKPDGIIGHSLGEVACGYADGCLSQREAVLAAYWRGQCIKDAHL<br>PPGSMAAVGLSWEECKQRCPAGVVPACHNSEDTVTISGPQAAVNEFVEQLKQEGVFAKEVRTGGLAFHSYF<br>MEGIAPTLLQALKKVIREPRPRSARWLSTSIPEAQWQSSLARTSSAEYNVNNLVSPVLFQEALWHIPEHAVV<br>LEIAPHALLQAVLKRGVKSSCTIIPLMKRDHKDNLEFFLTNLGKVHLTGINVNPNALFPPVEFPAPRGTPLISP<br>HIKWDHSQTWDVPVAEDFPNGSSSSSATVYSIDASAEKKAVAHGDGDTQRDLVKAVAHILGIRDLAGINLDS<br>TLADLGLDSLMGVEVRQILEREHDLVLPMREVRQLTLRKLQEMSSKTDSATDTTLEHHHHHHHH |
| pAR127 (nrFAS, Strep- and His-tagged) | ATGAGCGCTTGGAGCCATCCACAATTTGAGAAGGGTGGAGGTTCTGGCGGTGGATCGGGAGGTTCAGCGTG<br>GAGCCACCCGCAGTTCGAAAAAGGCGCCGGATCCGAGGAGGTGGTGATAGCCGGTATGTCGGGGAAGTTGC<br>CCGAGTCAGAGAACCTACAGGAGTTCTGGGCCAACCTCATTGGTGGTGTGGACATGGTCACAGATGATGAC<br>AGGAGATGGAAGGCTGGGCTCTATGGATTACCCAAGCGGTCTGGAAAGCTGAAGGATCTCTCCAAGTTCGA<br>CGCCTCCTTTTTTGGGGTCCACCCCAAGCAGGCACACACAATGGACCCCCAGCTTCGGCTGCTGTTGGAAGT<br>CAGCTATGAAGCAATTGTGGATGGAGGTATCAACCCAGCCTCACTCCGAGGAACGAACACTGGCGTCTGGG<br>TGGGTGTGAGTGGTTCAGAGGCATCCGAGGCCCTTAGCAGAGATCCCGAGACGCTTCTGGGCTACAGCATG<br>GTGGGCTGCCAGCGTGCAATGATGGCCAACCGGCTCTCTTTCTTCTTCGACTTCAAAGGACCAAGCATTGCC<br>CTGGACACAGCCTGCTCCTCCAGCTTGCTGGCACTACAGAATGCCTACCAGGCCATCCGTAGTGGGGAATGC<br>CCCGCGGCCCTTGTGGGTGGGATCAACCTGCTCCTGAAGCCGAACACCTCTGTGCAGTTCATGAAGCTGGGC<br>ATGCTCAGCCCGGACGGCACCTGCAGATCCTTTGATGATTCAGGGAGTGGATATTGTCGCTCTGAGGCTGT<br>TGTAGCAGTTCTGCTGACTAAGAAGTCCCTGGCTCGGCGGGTCTATGCCACGATTCTGAATGCCGGCACCAA<br>TACAGATGGCAGCAAGGAGCAAGGTGTAACATTCCCCTCTGGAGAAGTCCAAGAACAACTCATCTGCTCTC<br>TGTATCAGCCAGCTGGTCTGGCCCCGGAGTCGCTTGAGTATATTGAAGCCCATGGCACGGGCACCAAGGTG<br>GGTGACCCCCAGGAACTGAATGGCATTACTCGGTCCCTGTGCGCCTTCCGCCAGGCCCCTCTGTTAATTGGC<br>TCCACCAAATCCAACATGGGACACCCTGAGCCTGCCTCTGGGCTTGCAGCCCTGACCAAGGTGCTGTTATCC<br>CTGGAGCATGGGGTCTGGGCCCCTAACCTGCACTTCCACAACCCCAACCCTGAGATCCCAGCACTTCTTGAT<br>GGGCGGCTGCAGGTGGTCGATAGGCCCCTGCCTGTTCGTGGTGGCAACGTGGGCATCAACTCATTTGGCTTC<br>GGAGGCTCCAATGTTCATGTCATCCTCCAGCCCAACACACGGCAGGCCCCTGCGCCCACTGCACACGCTGCC<br>CTTCCCCATTTGCTGCACGCCAGTGGACGCACCTTAGAGGCAGTGCAGGACCTGCTGGAACAGGGCCGCCAG<br>CACAGCCAGGACCTGGCCTTTGTGAGCATGCTCAATGACATTGCGGCAACCCCTACAGCAGCCATGCCCTTC<br>AGGGGTTACACTGTGCTAGGTGTTGAGGGCCGTGTCCAAGAAGTGCAGCAAGTGTCCACCAACAAGCGCCC<br>ACTCTGGTTCATCTGCTCAGGGATGGGCACGCAGTGGCGCGGGATGGGGCTGAGCCTCATGCGCCTGGACA<br>GCTTCCGTGAGTCTATCCTGCGCTCCGATGAGGCTGTGAAGCCGTTGGGAGTGAAAGTGTCAGATCTGCTG<br>TTGAGCACAGATGAGCGCACCTTTGATGACATCGTGCATGCCTTTGTGAGCCTCACTGCCATCCAGATTGCC<br>CTCATCGACCTACTGACTTCTGTGGGACTGAAACCTGACGGCATCATTGGGCACTCCTTGGGAGAGGTTGCC<br>TGTGGCTATGCAGATGGCTGTCTCTCCCAGAGAGAGGCTGTGCTTGCAGCTTACTGGCGAGGCCAGTGCAT<br>CAAAGATGCCCACCTCCCGCCTGGATCCATGGCAGCTGTTGGTTTGTCCTGGGAGGAATGTAAACAGCGCTG<br>CCCCGCTGGCGTGGTGCCTGCCTGCCACAACTCTGAGGACACCGTGACCATCTCTGGACCTCAGGCTGCAGT<br>GAATGAATTTGTGGAGCAGCTAAAGCAAGAAGGTGTGTTTGCCAAGGAGGTACGAACAGGAGGCCTGGCTT<br>TCCACTCCTACTTCATGGAAGGAATTGCCCCCACATTGCTGCAGGCTCTCAAGAAGGTGATCCGGGAACCAC<br>GGCCGCGCTCGGCTCGATGGCTCAGCACCTCTATCCCTGAGGCCCAGTGGCAGAGCAGCCTGGCCCGCACAT<br>CTTCTGCCGAGTACAATGTCAACAACCTGGTGAGCCCTGTGCTCTTCCAGGAAGCACTGTGGCACATCCCTG<br>AGCATGCCGTGGTGCTGGAGATTGCGCCCCACGCACTGTTGCAGGCTGTCCTGAAGCGAGGCGTGAAGTCC<br>AGCTGCACCATCATTCCCTTGATGAAGAGGGATCATAAAGATAACTTGGAGTTCTTTCTCACCAACCTTGG<br>CAAGGTGCACCTCACAGGCATCAATGTCAACCCTAACGCCTTGTTCCCACCTGTGGAGTTCCCGGCTCCCCG<br>AGGGACTCCTCTCATCTCCCCTCACATCAAGTGGGACCACAGTCAGACTTGGGATGTCCCGGTTGCTGAGGA<br>CTTCCCAAACGGCTCCAGCTCCTCCTCTGCTACAGTCTACAGCATCGACGCCAGTGCAGAGAAGAAAGCTGT<br>GGCCCATGGGGACGGGGACACCCAGAGGGATCTGGTGAAAGCTGTAGCACACATCCTAGGCATCCGAGACC<br>TCGCAGGTATTAACCTGGACAGCACGCTGGCAGACCTCGGCCTGGACTCGCTCATGGGTGTGGAAGTTCGTC |

| Construct / Helix | Sequence from start to stop codon / as swapped |
|---|---|
| | AGATCCTGGAACGAGAACACGATCTGGTGCTGCCCATGCGTGAGGTGCGGCAGCTCACGCTGCGGAAACTT<br>CAGGAAATGTCCTCCAAGACTGACTCGGCTACTGACACGACACTCGAGCATCATCACCACCACCACCACCAC |
| DmbS Helix | VDDNRIQSNIATMRAMRLS |
| | GTCGACGACAACAGAATACAGAGCAACATAGCTACCATGCGAGCTATGAGGCTCTCT |
| TenS Helix | VDDTKVQMSLGTTRVMSVS |
| | GTTGACGACACCAAGGTGCAGATGAGCCTAGGTACCACGCGAGTCATGAGTGTCTCT |
| *E. coli* FabG Helix | ALSDDQRAGILAQVPAGRLGG |
| | GCGCTGAGCGATGACCAGCGTGCGGGTATCCTGGCGCAGGTTCCTGCGGGTCGCCTCGGCGGC |
| *H. pylori* FabG Helix | NLKDELKADYVKNIPLNRLGS |
| | AATTTGAAAGACGAACTCAAAGCGGATTATGTTAAAAACATTCCTTTAAACAGGCTAGGGTCT |
